# Distinct roles of two thalamostriatal systems in learning processes of visual discrimination in common marmosets

**DOI:** 10.1101/2024.06.06.597696

**Authors:** Shigeki Kato, Masateru Sugawara, Miwako Yamasaki, Masahiko Watanabe, Ken-ichi Inoue, Katsuki Nakamura, Daisuke Koketsu, Satomi Chiken, Atsushi Nambu, Masahiko Takada, Kazuto Kobayashi

## Abstract

The thalamostriatal projections arising from the intralaminar thalamic nuclei (ILN) constitute the principal source of input information to specified subregions of the striatum, a key structure of the cortico-basal ganglia circuitry. However, the roles of primate ILN in cortico-basal ganglia circuit functions remain unclear. Here, we performed immunotoxin-induced selective targeting of two representative structures of the ILN, the parafascicular nucleus (Pf) and centre médian nucleus (CM) projecting to the caudate nucleus (Cd) and putamen (Pu), respectively, in common marmosets. Elimination of Pf-Cd neurons resulted in impaired reversal learning of a two-choice visual discrimination task, whereas removal of CM-Pu neurons disturbed the task acquisition. No marked impact of such manipulations was observed on either motor skill learning or spontaneous locomotor activity. Our findings reveal that the two thalamostriatal systems play distinct roles in the learning processes of external cue-dependent decision-making in nonhuman primates.

## Introduction

The striatum is a key structure of the cortico-basal ganglia circuitry, which controls a variety of behaviors including motor control, motor learning, instrumental learning, and behavioral switching^1–5^. It is divided into the caudate nucleus (Cd) and putamen (Pu), each of which consists of multiple sectors with functional diversity in primates^6,7^. The intralaminar thalamic nuclei (ILN), which comprise the centre médian nucleus (CM) and parafascicular nucleus (Pf) in primates, constitute the principal source of afferent fibers projecting to specified striatal subregions^8,9^. The CM/Pf neurons in macaque monkeys are reportedly involved in the attention process for detection of an unpredictable stimulus or are associated with orientation to sensory events^10,11^. The CM neurons also seem to be engaged in the regulation of a response bias toward a reward amount^12^. However, specific behavioral roles of the projections from the CM/Pf to striatal target regions have not yet been defined.

Parkinson’s disease (PD) is characterized by the marked loss of dopamine neurons from the substantia nigra pars compacta (SNc), in which neurodegeneration is caused by complex mechanisms such as alpha-synuclein accumulation, oxidative stress, mitochondrial dysfunction, and cellular calcium imbalance^13,14^. The major symptoms in PD are prominent motor disturbances, which are accompanied by various cognitive impairments including deficits in learning processes and behavioral flexibility^15–18^. In addition to SNc dopamine neurons, neurodegeneration in PD also occurs in other brain regions^19–21^. Postmortem brain analysis has shown that neurons in the CM/Pf of ILN display extensive degeneration in PD patients^22–24^. Moreover, the number of CM/Pf neurons and density of their striatal terminals are reduced in parkinsonian monkeys treated with 1-methyl-4-phenyl-1,2,3,6-tetrahydropyridine (MPTP)^25,26^. These observations suggest that neuronal loss in the CM/Pf may be involved, at least in part, in motor and/or cognitive impairments in PD, although its pathogenesis remains unclear.

In the present study, we studied the possible involvement of the two representative thalamostriatal systems in common marmosets, focusing on Pf projection to the Cd and CM projection to the Pu, in motor and cognitive functions. For this purpose, we applied an immunotoxin-induced pathway-selective targeting technique, which has been used for the analysis of thalamostriatal functions in rodents^27,28^. Our results demonstrate that the Pf-Cd neurons in marmosets play an essential role in reversal learning of a visual discrimination task, whereas the CM-Pu neurons contribute to acquisition of the task. On the other hand, these two neuronal populations appear to exert no marked influence on either motor skill learning or spontaneous locomotor activity. Thus, the present findings reveal that the Pf-Cd and CM-Pu thalamostriatal systems provide distinct roles in the learning processes of external cue-dependent decision-making, suggesting that neuronal loss in the CM/Pf may represent a certain aspect of cognitive impairment in PD.

## Results

### Thalamostriatal projections in common marmosets

Previous anatomical studies have indicated that Pf and CM neurons mainly innervate the Cd and Pu, respectively, in the striatum of macaque monkeys^8^. To confirm the patterns of these thalamostriatal projections in common marmosets, we carried out retrograde labeling of neural pathways using a lentiviral vector with neuron-specific retrograde gene transfer (NeuRet)^29,30^ (see Fig. 1a for the experimental strategy). One marmoset was administered unilateral injections of the NeuRet vector encoding red fluorescent protein (RFP) (3.36 × 10^12^ genome copies/mL) into the Cd, mainly the head part (1.0 µL/site, 6 sites); and the NeuRet vector encoding green fluorescent protein (GFP) (3.27 × 10^12^ genome copies/mL) was injected into the Pu, particularly the anteroposterior middle region (1.0 µL/site, 4 sites) (see Fig. 1b for the injection sites shown on magnetic resonance [MR] images), because these regions have dense connections with some associative cortical areas^31–33^. Three weeks after the injections, sections through the ILN were prepared and stained with anti-RFP and anti-GFP antibodies. A number of RFP immunopositive signals were visualized in the Pf subregion surrounding the fasciculus retroflexus, and many GFP-positive signals were detected in the CM subregion localized laterally from the Pf (see a typical image in Fig. 1c). Magnified views of the image showed the presence of RFP and GFP signals in neurons in the two ILN subregions (Fig. 1d). These data indicate that the main connections of Pf neurons to the Cd and CM neurons to the Pu in common marmosets are consistent with the projection patterns of thalamostriatal systems in macaque monkeys^8^.

**Fig. 1.**
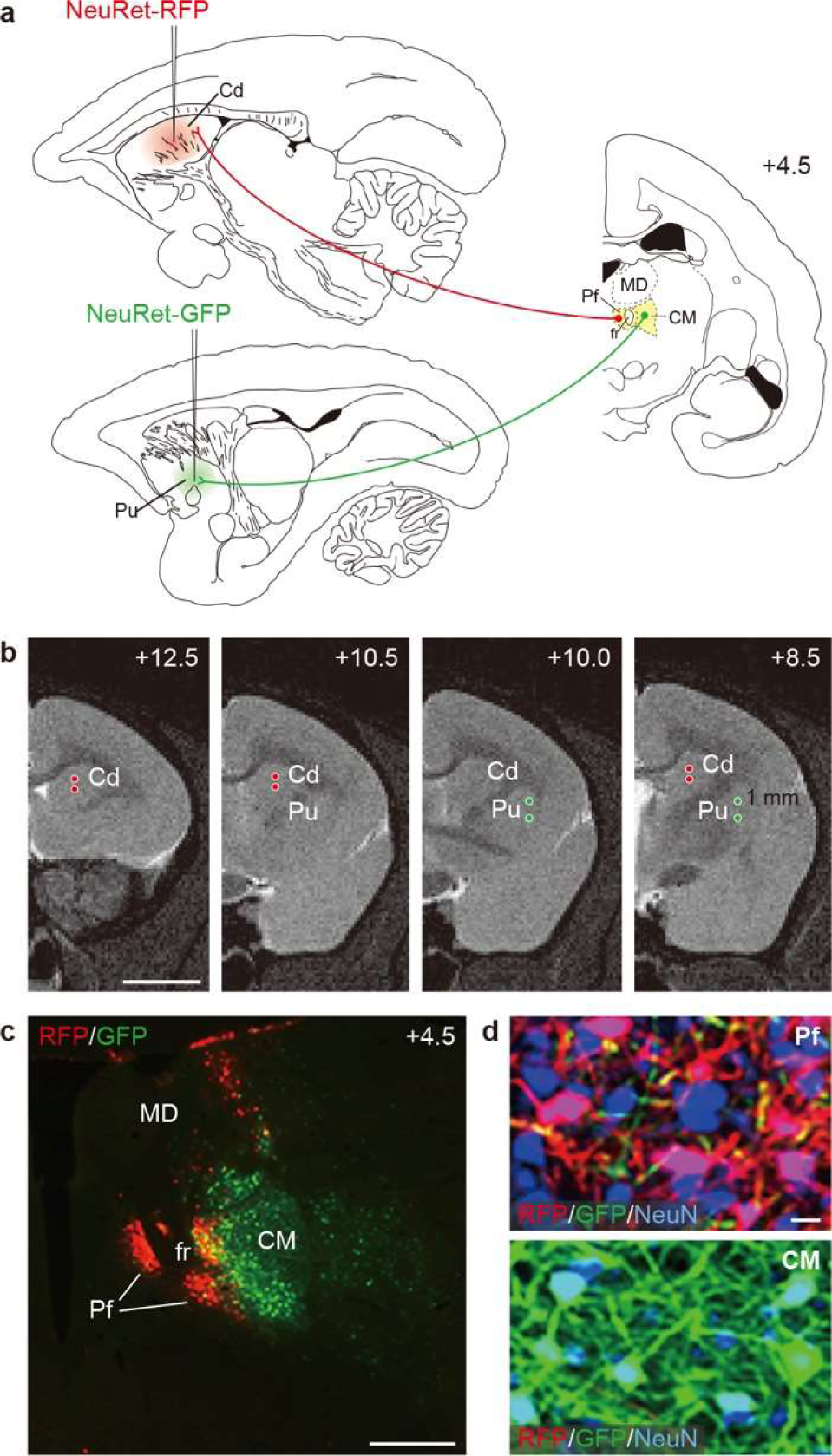
Projection patterns of two thalamostriatal systems in common marmosets. **a** Schematic illustration of an experimental strategy for retrograde labeling of neural pathways. NeuRet-RFP and NeuRet-GFP vectors were injected into the Cd (mainly its head part) or Pu (especially its anteroposterior middle part) of the common marmoset MAR-0, and sections through the ILN were immunostained for transgene detection. **b** Coordinates for the vector injection. Red and green circles indicate the injection sites into the Cd and Pu, respectively, on a series of MR images through a coronal plane. **c** Double immunostaining for RFP and GFP. A typical image of CM/Pf complex sections is shown. **d** Magnified views of the image for the Pf (upper) and CM (lower) subregions. NeuN was co-stained as a marker for neurons. The anteroposterior coordinates (mm) from bregma are shown. fr, fasciclus retroflexus. MD, mediodorsal thalamus. Scale bars: 5 mm (**b**), 1 mm (**c**), 20 mm (**d**).

### Immunotoxin-induced pathway-selective targeting of Pf-Cd neurons

We removed the Pf-Cd system using immunotoxin-induced pathway-selective targeting^27,28^ (see Fig. 2a for the experimental strategy). The NeuRet vector encoding a chimeric gene composed of human interleukin-2 receptor α-subunit (IL-2Rα) fused to GFP (3.55 × 10^12^ genome copies/mL) was bilaterally injected into the Cd head corresponding to the site examined for the above tracing experiments (1.0 µL/site, 6 sites/hemisphere) (see Fig. 2b for the coronal planes 1–3 on a sagittal MR image; and Fig. 2c for the injection sites on three coronal images, which were obtained from a representative injected animal). Three weeks later, a solution containing recombinant immunotoxin (ITX) (25 ng/µL), which specifically recognizes human IL-2Rα and induces the apoptosis of cells expressing the receptor^27^, was bilaterally injected into the Pf (0.5 µL/site, 1 site/hemisphere) (see Fig. 2b for coronal plane 4 on the sagittal MR image; and Fig. 2d for the injection sites on the coronal image from the same animal). For the controls, phosphate-buffered saline (PBS) was injected into the Pf of the vector-injected animals. The coordinates used for injections of the viral vector into the Cd head and ITX/PBS into the Pf for each animal (*n* = 4 for the ITX-injected group and *n* = 3 for the PBS-injected group) are summarized in Supplementary Table 1.

**Fig. 2.**
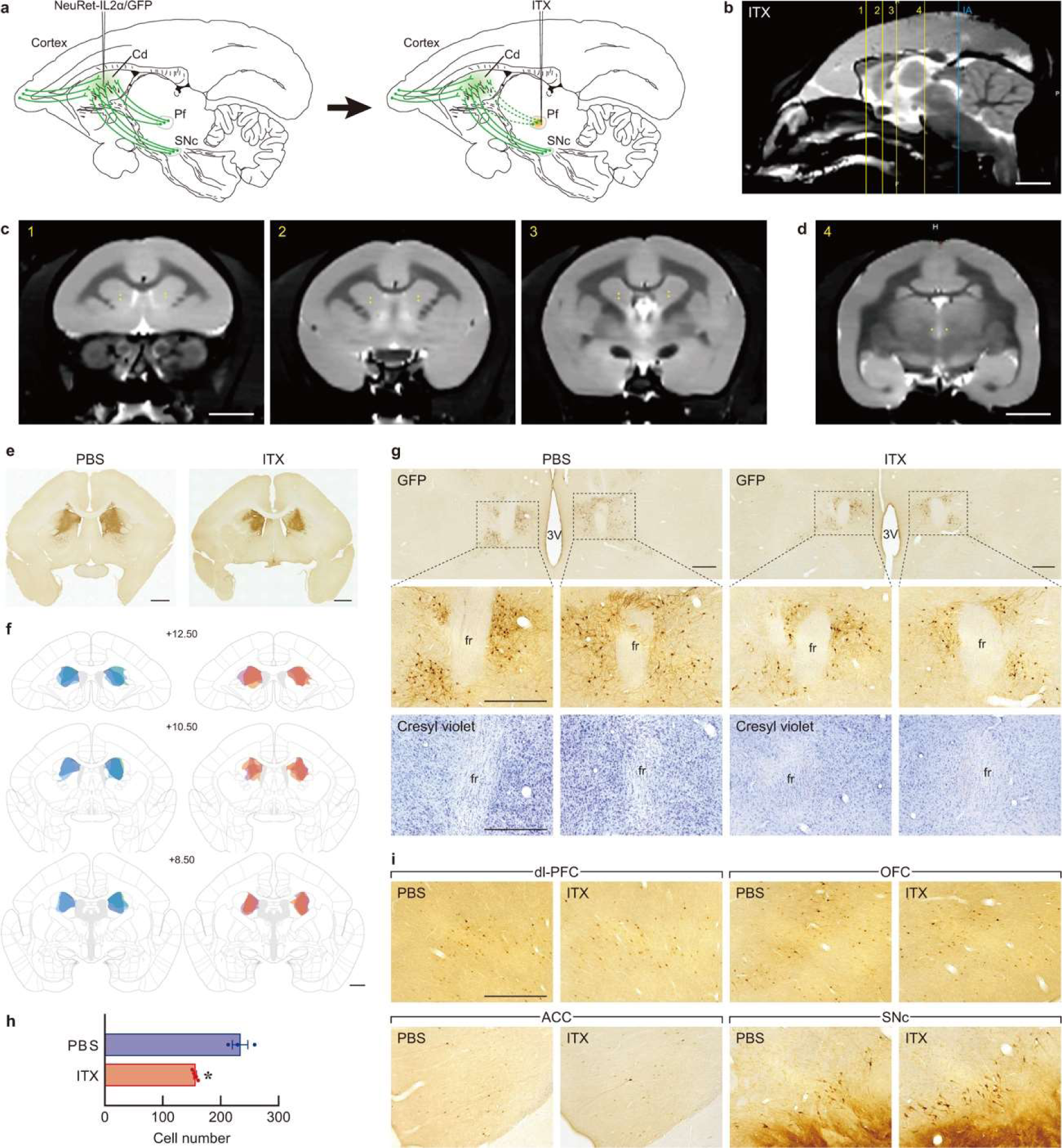
Selective removal of Pf-Cd neurons in common marmosets. **a** Experimental strategy for targeting the Pf-Cd system. The marmosets received bilateral injections of NeuRet-IL2Rα/GFP vector into the Cd, mainly the Cd head and then ITX solution into the Pf, resulting in the elimination of Pf-Cd neurons. **b-d** Representative MR images used for injections of the viral vector and ITX/PBS obtained from the marmoset MAR-6. Sagittal MR image showing the coronal planes 1–3 for vector injection and plane 4 for ITX injection (**b**). Coordinates for vector injection on three coronal MR images (**c**). Coordinates for ITX injection on a coronal MR image, in which injections sites are indicated by yellow spots (**d**). **e, f** Immunohistochemical staining of sections through the striatum of the IT/PBS-injected marmosets. Typical microscopic images (**e**) and schematic illustration summarizing the distribution of GFP-positive terminals in individual animals (**f**). **g** Immunostaining of sections through the ILN of the IT/PBS-injected groups. Middle images are magnified views of the rectangles in the upper images. Cresyl violet staining of the sections are shown in the lower images. The typical microscopic images of ITX- and PBS-injected groups are shown. 3V, third ventricle; fr, fasciculus retroflexus. **h** Cell counts of GFP-positive cells in the Pf. **i** GFP staining of sections through the cerebral cortical areas and ventral midbrain of the ITX/PBS-injected groups. The typical photos of the two groups were taken from the same animals as above. ACC, anterior cingulate cortex; dl-PFC, dorsolateral prefrontal cortex; OFC, orbitofrontal cortex; and SNc, substantia nigra pars compacta. *n* = 3 for the PBS-injected group; and *n* = 4 for the ITX-injected group. \*\**p* < 0.01 (unpaired *t*-test). Data are presented as the mean ± SEM. The anteroposterior coordinates (mm) from bregma are shown. Scale bars: 5 mm (**b**-**d**), 2 mm (**e**,**f**), 500 mm (**g**,**i**).

First, behavioral tests were conducted (see below for the results; Figs. 3 and 4). Then the animals were sacrificed followed by histological analyses (Fig. 2e–i). Coronal sections through the striatum, ILN, and cerebral cortical areas were prepared and stained with anti-GFP antibody. A large number of GFP-positive nerve terminals were widely localized in the Cd in both ITX- and PBS-injected groups (see Fig. 2e for typical microscopic images of the Cd sections, and Fig. 2f for the distribution of immunopositive terminals in the striatum from individual animals). Although many GFP-positive cells were detected in the Pf subregion in the PBS-injected group, the number of cells in the corresponding subregion was reduced in the ITX-injected group (Fig. 2g, upper and middle images), showing a significant reduction in cell number to 66.7% in the ITX-injected group (155.83 ± 2.27/section) compared to the control group (233.47 ± 13.35/section; unpaired *t*-test, *t*_5_ = 6.758, *p* = 0.001) (Fig. 2h). Cresyl violet staining of Pf sections excluded nonspecific injury of the tissues after ITX treatment (Fig. 2g, lower images). In addition, there were no apparent changes in the distribution of immunopositive cells in some cortical areas, including the dorsolateral prefrontal cortex, orbitofrontal cortex, anterior cingulate cortex, and SNc in the ventral midbrain between the ITX- and PBS-injected groups (Fig. 2i), suggesting no damage to other striatal inputs into the Cd head with the exception of Pf-Cd neurons.

**Fig. 3.**
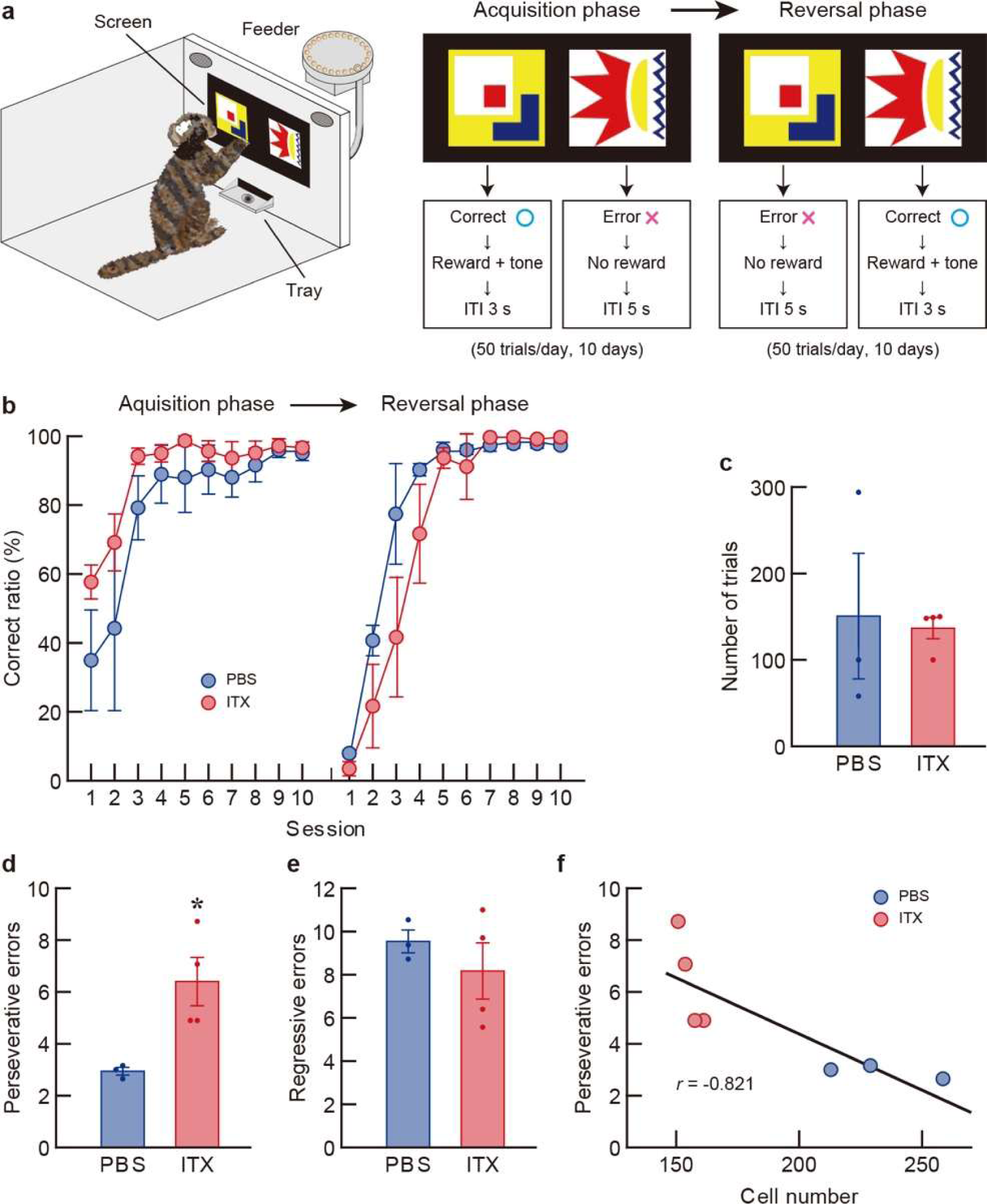
Acquisition and reversal learning of a two-choice visual discrimination task in common marmosets lacking Pf-Cd neurons. **a** Schematic diagram of the visual discrimination task. Each trial started by touching a red square image at the center of the front screen. After 1 s, two graphic images were presented on both sides of the screen, and the locations of the images were randomly changed in each trial. The marmosets were required to touch one of the images (correct image), resulting in a solid reward through the food tray from the feeder with a tone presentation. The animals were trained 50 trials in each daily session for 10 consecutive days during the acquisition phase. Then the correct image was converted to another one, and they were trained 50 trials in each daily session for 10 consecutive days again during the reversal phase. **b** Learning curves in the acquisition and reversal phases. The correct ratio in each session was plotted along with the progress of training. **c** Number of trials required to reach a criterion of 75% correction in the acquisition phase. **d, e** Numbers of perseverative and regressive errors in the reversal phase. **f** Correlation between perseverative error number and Pf cell number in individual animals. Pearson’s correlation coefficient: *r* = -0.821, *p* = 0.024. *n* = 4 for the ITX-injected group, and *n* = 3 for the PBS-injected group. \**p* < 0.05 (unpaired *t*-test). Data are presented as the mean ± SEM.

**Fig. 4.**
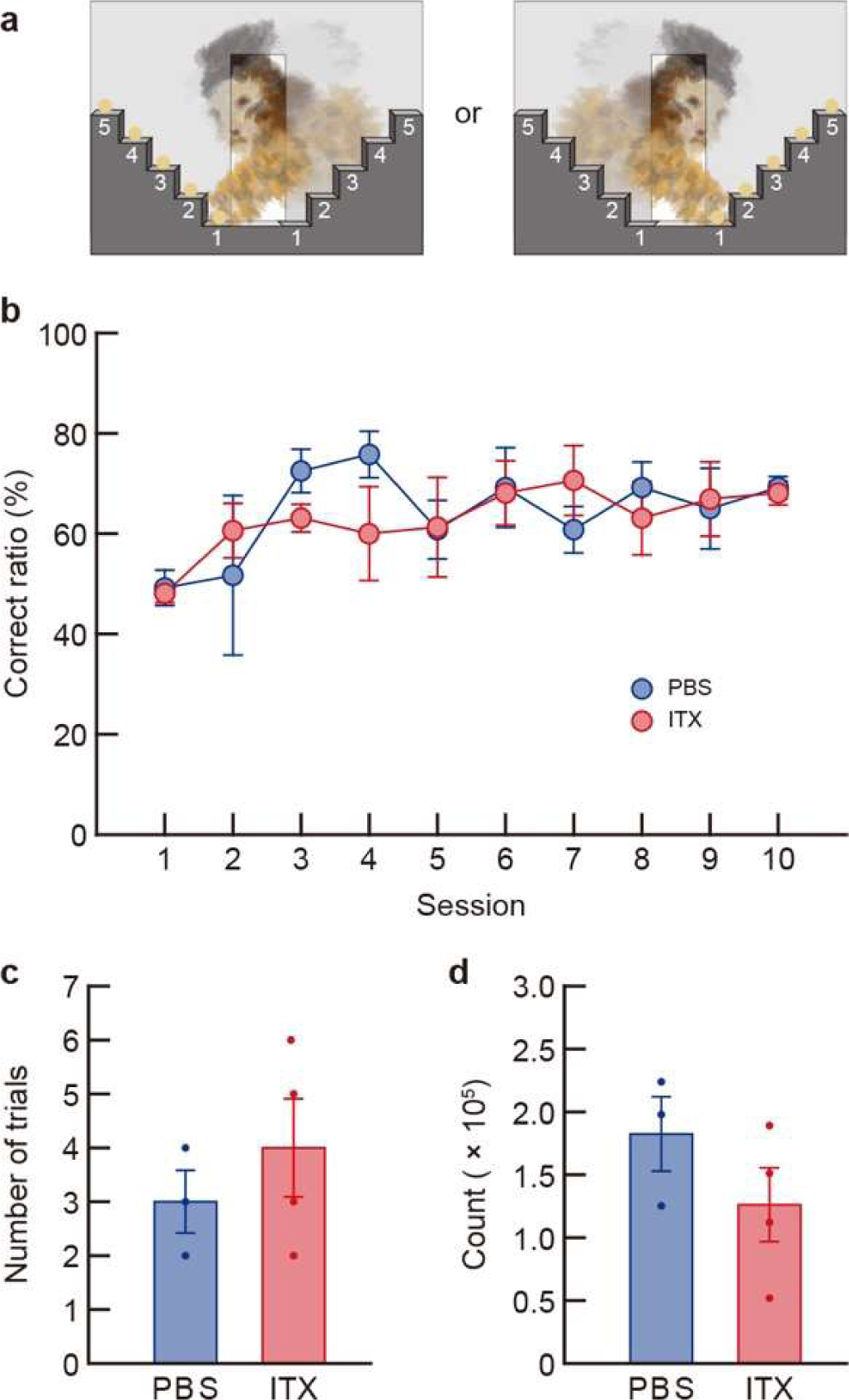
Motor skill learning and spontaneous locomotor activity in common marmosets lacking Pf-Cd neurons. **a** Schematic diagram of the staircase foraging task. Five solid rewards were placed on either of the left or right side of the staircase apparatus. The marmosets were required to reach and retrieve the reward through the central slit by using a unilateral arm opposite to the side of the stairs. In each daily session, they were trained every 5 trials on either side and then on another side. The session continued for 10 consecutive days. **b** Learning curves in the acquisition of motor skills. The correct ratio in each session was plotted along with the progress of training. **c** Number of trials required to reach a criterion of 70% correction. **d** Locomotion activity counts. The counts were measured in a home cage for a 12 h-test period. *n* = 4 for the ITX-injected group, and *n* = 3 for the PBS-injected group. Data are presented as the mean ± SEM.

### Targeting of Pf-Cd neurons impairs reversal learning of visual discrimination

Common marmosets that received injections with the viral vector and ITX/PBS were used for the analyses of behavioral tasks that require the functions of cortico-basal ganglia circuit. To investigate the roles of the Pf-Cd system in the acquisition of external cue-dependent discrimination learning and its reversal of the learned behavior, we conducted a two-choice visual discrimination task^34,35^. In this task, after animals learned to touch a single graphic image presented on a front screen during the habituation phase, two graphic images were displayed simultaneously on the left and right sides of the screen. Animals learned to touch one of the two images, which was associated with a reward (see Fig. 3a). Two graphic images were selected from a series of images (Supplementary Fig. 1), and the locations of the paired images were randomly shifted in each trial. The acquisition of visual discrimination and its reversal performance were tested. The ITX- and PBS-injected groups learned normally single image touches during the habituation phase (Supplementary Fig. 2). The correct ratio was increased during the acquisition phase in a similar pattern between the ITX- and PBS-injected groups (two-way analysis of variance [ANOVA], group, *F* _(1, 5)_ = 2.491, *p* = 0.175; session, *F* _(9, 45)_ = 16.190, *p* < 0.001; interaction, *F* _(9, 45)_ = 0.920, *p* = 0.517) (Fig. 3b), and the numbers of trials, which reached a criterion of 75% correction, were comparable between the two injected groups (136.75 ± 12.26 and 150.67 ± 72.69 for the ITX- and PBS-injected groups, respectively; unpaired *t*-test, *t*_5_ = 0.189, *p* = 0.867) (Fig. 3c). During the reversal phase, the increase in the correct ratio appeared to be modestly slower in the early phase, although there was no significant difference between the groups (two-way ANOVA, group, *F* _(1, 5)_ = 2.954, *p* = 0.146; session, *F* _(9, 45)_ = 41.280, *p* < 0.001; interaction, *F* _(9, 45)_ = 1.510, *p* = 0.174) (Fig. 3b). Error responses in the reversal phase were evaluated as “perseverative errors,” defined as error responses that occur until an animal first touches the new correct image; and “regressive errors,” defined as error responses that occur afterwards^36,37^. The number of perseverative errors was significantly larger in the ITX-injected group (6.40 ± 0.93) compared to the PBS-injected group (2.94 ± 0.15) (unpaired *t*-test, *t*_5_ = 3.681, *p* = 0.032) (Fig. 3d), whereas the number of regressive errors was similar between the two groups (8.17 ± 1.30 and 9.54 ± 0.53 for the ITX- and PBS-injected groups, respectively; unpaired *t*-test, *t*_5_ = 0.982, *p* = 0.383) (Fig. 3e). In addition, the number of perseverative errors in both groups was negatively correlated with the cell number in the Pf subregion, which was obtained from immunostaining as shown in Fig. 2h (Pearson’s correlation coefficient: *r* = -0.821, *p* = 0.024) (Fig. 3f). These results demonstrate that the elimination of Pf-Cd neurons in the marmosets resulted in impaired reversal learning of the visual discrimination task, but had no effect on acquisition of the same task, suggesting the importance of these neurons in the flexible switching of learned behavior.

Next, to study the functions of Pf-Cd neurons in motor skill learning, we employed a staircase foraging task^38,39^. In this task, animals learned to reach and retrieve solid rewards through a central plate slit by using a left or right arm opposite the side of the stairs of the apparatus consisting of five stairs on each side (see Fig. 4a). The correct ratio appeared to gradually increase along with the progress of sessions in both the ITX- and PBS-injected groups, showing no significant difference between the two injected groups (two-way ANOVA, group, *F* _(1, 5)_ = 0.037, *p* = 0.856; session, *F* _(9, 45)_ = 3.218, *p* = 0.004; interaction, *F* _(9, 45)_ = 1.116, *p* = 0.371) (Fig. 4b). There was also no significant difference in the number of trials required to reach a criterion of 70% correction between the two groups (4.00 ± 0.91 and 3.00 ± 0.58 for the ITX- and PBS-injected groups, respectively; unpaired *t*-test, *t*_5_ = 0.845, *p* = 0.437) (Fig. 4c). In addition, we assessed the spontaneous locomotion of the marmosets lacking Pf-Cd neurons by using a miniatured accelerometer^40,41^. Locomotion activity was counted in a home cage for a test period of 12 h. The counts were not significantly different between the ITX- and PBS-injected groups (1.26 ± 0.29 × 10^5^ and 1.82 ± 0.30 × 10^5^, respectively; unpaired *t*-test, *t*_5_ = 1.322, *p* = 0.243) (Fig. 4d). These data indicate that Pf-Cd neurons did not seem to be implicated in the acquisition of motor skills or the control of spontaneous locomotion activity in the marmosets.

### Immunotoxin-induced pathway-selective targeting of CM-Pu neurons

For elimination of the CM-Pu system in common marmosets (see Fig. 5a), the NeuRet-IL-2Rα/GFP vector (3.80 × 10^12^ genome copies/mL) was bilaterally injected into the middle Pu, corresponding to the site examined for the above tracing experiments (1.0 µL/site, 4 sites/hemisphere) (see Fig. 5b for the coronal planes 1/2 on a sagittal MR image; and Fig. 5c for the injection sites on two coronal images, which were obtained from a representative injected animal). Three weeks later, ITX solution (25 ng/µL) or PBS was bilaterally injected into the CM (0.5 µL/site, 2 sites/hemisphere) (see Fig. 5b for the coronal plane 3/4 on the sagittal MR image; and Fig. 5d for the injection sites on the coronal images from the same animal). The coordinates used for injections of the viral vector into the middle Pu and ITX/PBS into the CM for each animal (*n* = 4 for each group) are shown in Supplementary Table 2.

**Fig. 5.**
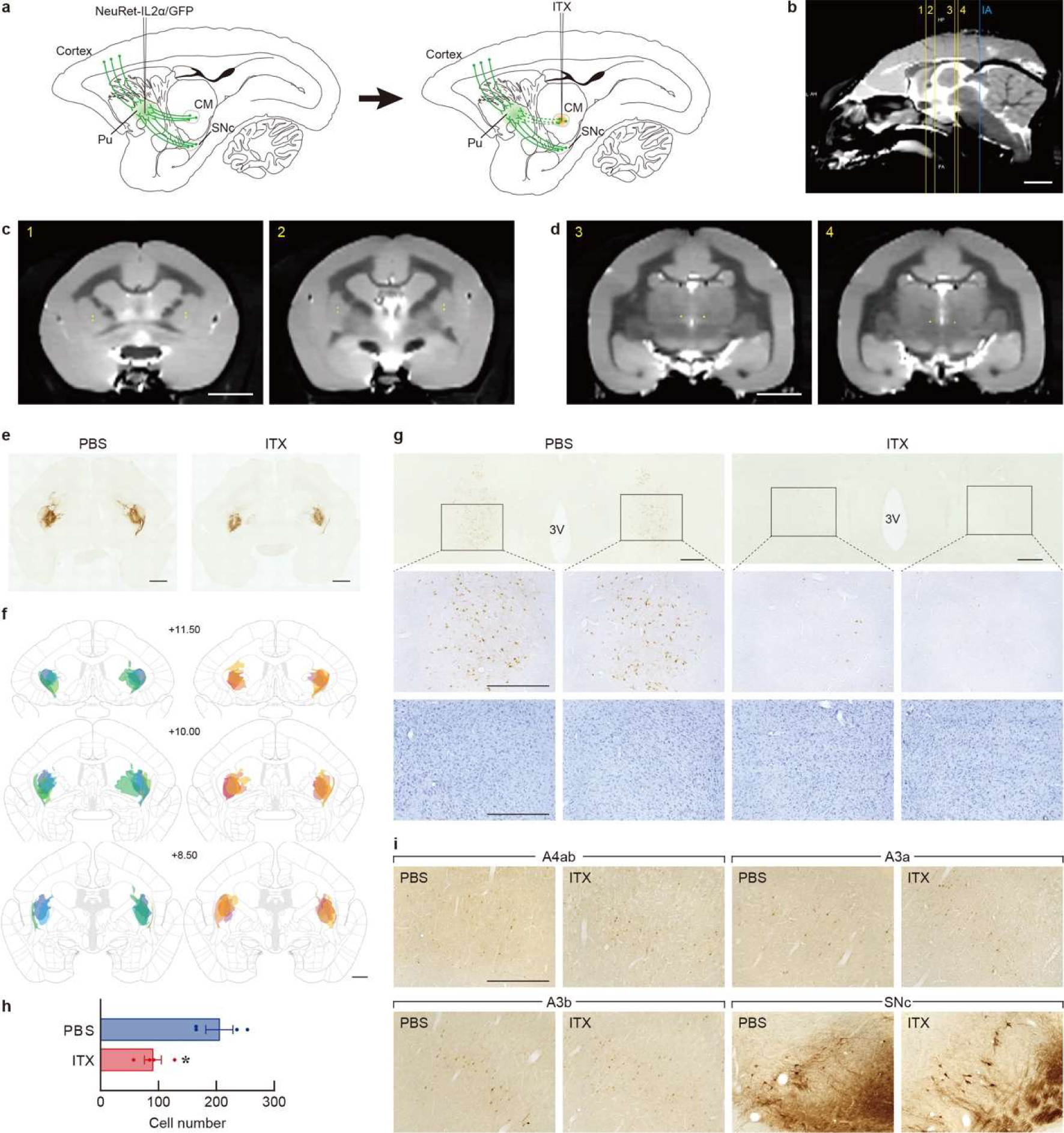
Selective elimination of CM-Pu neurons in common marmosets. **a** Experimental strategy for targeting the CM-Pu system. The marmosets received bilateral injections of NeuRet-IL2Rα/GFP vector into the Pu, especially the middle Pu and then ITX solution into the CM, leading to the ablation of CM-Pu neurons. **b-d** Representative MR images used for treatments of the viral vector and ITX/PBS taken from the marmoset MAR-12. Sagittal MR image showing the coronal planes 1/2 for vector injection and the planes 3/4 for ITX injection (**b**). Coordinates for vector injection on two coronal MR images (**c**). Coordinates for ITX injection on two coronal MR images, in which injections sites are shown by yellow spots (**d**). **e, f** Immunohistochemistry of sections through the striatum of the IT/PBS-injected marmosets. Typical microscopic images (**e**) and schematic representation summarizing the distribution of immuno-positive terminals in individual animals (**f**). **g** Immunostained sections through the ILN of the ITX/PBS-injected groups. Middle images are magnified views of the rectangles in the upper images. Cresyl violet staining of the sections are indicated in the lower images. The typical microscopic images of the ITX- and PBS-injected groups are presented. 3V, third ventricle. **h** Cell counts of immunopositive cells in the CM. **i** GFP staining of sections through the cortical areas and ventral midbrain of the ITX/PBS-injected groups. Representative photos of the two groups were obtained from the same animals as above. A4ab, anterior cingulate cortex; A3a, dorsolateral prefrontal cortex; A3b, orbitofrontal cortex; and SNc, substantia nigra pars compacta. *n* = 4 for each group. \*\**p* < 0.01 (unpaired *t*-test). Data are presented as the mean ± SEM. The anteroposterior coordinates (mm) from bregma are shown. Scale bars: 5 mm (**b**-**d**), 2 mm (**e**,**f**), 500 mm (**g**,**i**).

After the behavioral tests (Figs. 6 and 7), histological analyses were performed (Fig. 5e–i). Sections of the brain regions used for the analysis of CM-Pu targeting were prepared and immunostained for GFP. A number of GFP-positive terminals were extensively visualized in the middle Pu in both the ITX- and PBS-injected groups (see Fig. 5e for typical microscopic images of the Pu sections, and Fig. 5f for the distribution of positive terminals in the striatum from individual animals). The number of cells in the CM subregion, compared to the PBS-injected group, was decreased in the ITX-injected groups (Fig. 5g, upper and middle images), indicating a significant decrease in cell number to 44.0% in the ITX-injected group (90.33 ± 14.59/section) compared to the control group (205.15 ± 23.49/section) (unpaired *t*-test, *t*_6_ = 4.152, *p* = 0.006) (Fig. 5h). Cresyl violet staining of CM sections ruled out nonspecific injury of the tissues after ITX treatment (Fig. 5g, lower images). There were no gross variations in the distribution of immunopositive cells in some cortical areas, including the anterior cingulate cortex (A4ab), dorsolateral prefrontal cortex (A3a), orbitofrontal cortex (A3b), and in the SNc between the injected groups (Fig. 5i), suggesting no damage to other striatal inputs into the middle Pu with the exception of CM-Pu neurons.

**Fig. 6.**
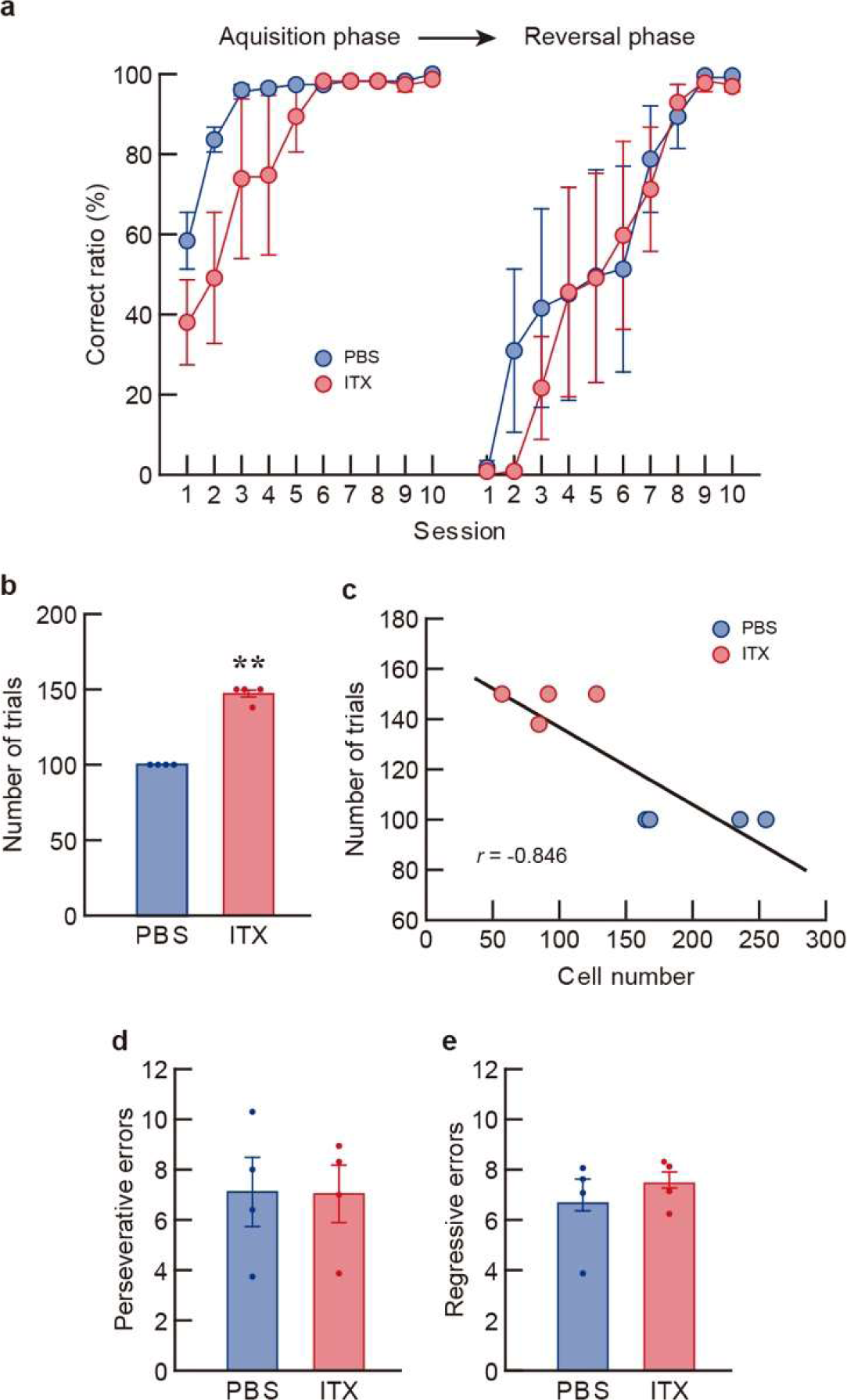
Acquisition and reversal learning of the visual discrimination task in common marmosets lacking CM-Pu neurons. **a** Learning curves in the acquisition and reversal phases. The correct ratio in each session was plotted in accordance with the progress of training. **b** Number of trials required to reach a 75% correction criterion in the acquisition phase. **c** Correlation between trial number for the acquisition criterion and CM cell number in individual animals. Pearson’s correlation coefficient: *r* = -0.846, *p* = 0.008. **d, e** Numbers of perseverative and regressive errors in the reversal phase. *n* = 4 for each group. \*\**p* < 0.01 (unpaired *t*-test). Data are presented as the mean ± SEM.

**Fig. 7.**
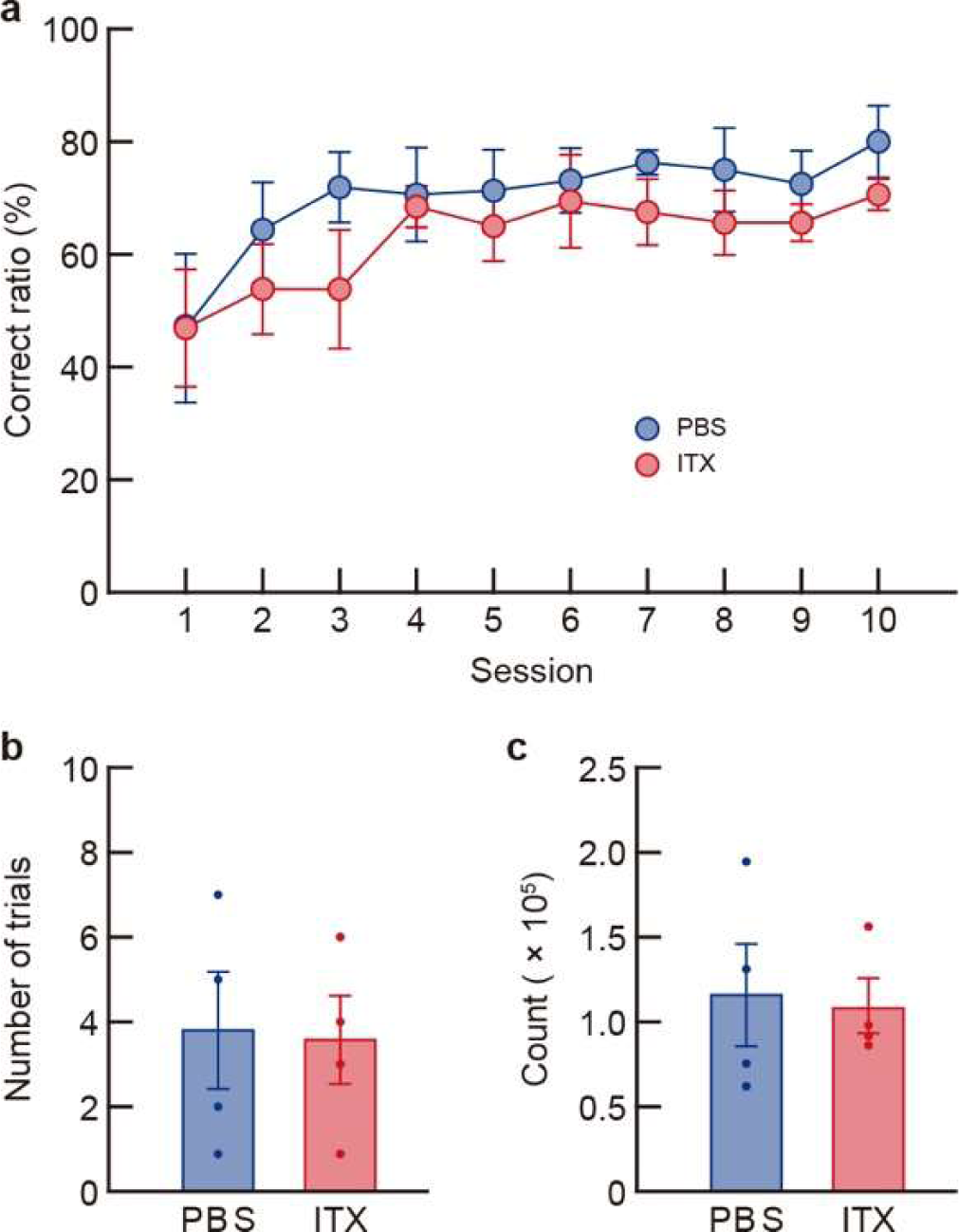
Motor skill learning and spontaneous locomotor activity in common marmosets lacking CM-Pu neurons. **a** Learning curves in the acquisition of motor skills. The correct ratio in each session was plotted in accordance with the progress of training. **b** Number of trials required to reach a 70% correction criterion. **c** Locomotion activity counts. The counts were monitored in a home cage for a 12 h-test period. *n* = 4 for each group. Data are presented as the mean ± SEM.

### Targeting of CM-Pu neurons impairs acquisition of visual discrimination

To study the behavioral roles of the CM-Pu system, we tested the same tasks as those used for analysis of the Pf-Cd system with the marmosets lacking CM-Pu neurons. When the two-choice visual discrimination task was conducted, the ITX- and PBS-injected groups showed normal single image touches during the habituation phase (Supplementary Fig. 3). The increase in the correct ratio during the acquisition phase appeared to be delayed in the ITX-injected group compared to the PBS-injected group (two-way ANOVA, group, *F* _(1, 6)_ = 1.840, *p* = 0.224; session, *F* _(9, 54)_ = 15.021, *p* < 0.001; interaction, *F* _(9, 54)_ = 2.080, *p* = 0.048) (Fig. 6a). The number of trials, which reached a criterion of 75% correction, was significantly elevated in the ITX-injected group (147.00 ± 3.00) compared to the control group (100.00 ± 0.00) (unpaired *t*-test, *t*_6_ = 15.667, *p* = 0.001) (Fig. 6b). The number of trials for the acquisition criterion was negatively correlated with the cell number in the CM subregion, which was obtained from immunostaining as shown in Fig. 5h (Pearson’s correlation coefficient: *r* = -0.846, *p* = 0.008) (Fig. 6c). During the reversal phase, the correct ratio was similarly increased between the two groups (two-way ANOVA, group, *F* _(1, 6)_ = 0.088, *p* = 0.777; session, *F* _(9, 54)_ = 16.326, *p* < 0.001; interaction, *F* _(9, 54)_ = 0.458, *p* = 0.896) (Fig. 6a). In addition, the numbers of perseverative errors did not differ between the two groups (7.03 ± 1.13 and 7.11 ± 1.38 for the ITX- and PBS-injected groups, respectively; unpaired *t*-test, *t*_6_ = 0.044, *p* = 0.966) (Fig. 6d), and the regressive error numbers were also similar between the groups (7.45 ± 0.48 and 6.66 ± 0.95 for the ITX- and PBS-injected groups, respectively; unpaired *t*-test, *t*_6_ = 0.752, *p* = 0.481) (Fig. 6e). These results demonstrate that the removal of CM-Pu neurons in the marmosets caused impaired acquisition of the visual discrimination task with no influence on the reversal of the same task, suggesting the significance of these neurons in the new learning of discrimination dependent on external cues.

For the staircase foraging task, the correct ratio exhibited a gradual increase in accordance with the session progress in both the ITX- and PBS-injected groups (two-way ANOVA, group, *F* _(1, 6)_ = 1.238, *p* = 0.308; session, *F* _(9, 54)_ = 4.047, *p* < 0.001; interaction, *F* _(9, 54)_ = 0.359, *p* = 0.950) (Fig. 7a), showing no significant difference in the number of trials required to reach a criterion of 70% correction between the two groups (3.50 ± 1.04 and 3.75 ± 1.38 for the ITX- and PBS-injected groups, respectively; unpaired *t*-test, *t*_6_ = 0.145, *p* = 0.890) (Fig. 7b). In addition, locomotion activity counts were not significantly different between the two injected groups (1.08 ± 0.16 × 10^5^ and 1.16 ± 0.30 × 10^5^ for the ITX- and PBS-injected groups, respectively; unpaired *t*-test, *t*_6_ = 0.226, *p* = 0.829) (Fig. 7c). The data indicate that CM-Pu neurons do not appear to be responsible for motor skill learning and spontaneous locomotion activity in marmosets.

## Discussion

We performed immunotoxin-induced pathway-selective targeting of the two major thalamostriatal systems, the Pf-Cd and CM-Pu projections, in common marmosets. Our findings provide evidence that Pf-Cd neurons play an important role in reversal learning of a two-choice visual discrimination task, whereas CM-Pu neurons contribute to acquisition of the task. This clearly indicates the distinct roles of these two thalamostriatal systems in the learning processes of external cue-dependent decision-making. The use of the pathway-selective targeting strategy has successfully revealed the specific involvement of Pf-Cd versus CM-Pu neurons in cognitive functions in nonhuman primates.

Previous electrophysiological recording and neuroimaging studies have shown that the Cd neurons are involved in different types of behavioral switching in humans and nonhuman primates^42–45^. The orbital and dorsolateral prefrontal cortical areas in macaque monkeys project mainly to the head or body part of the Cd^31^, and the corresponding areas in common marmosets have dense innervations to the head and body parts of the Cd, respectively^32^.The orbital frontal cortex is the critical locus in reversing the associated memory, and the lateral prefrontal cortex is essential for shifting attention between perceptual dimensions of complex stimuli^46,47^. In addition, neuroimaging investigations have shown that the Pu is recruited during the initial acquisition of instrumental learning in humans^48^. The parietal association cortices project to the middle Pu, in particular to its dorsal region^33^, although the presence of the corresponding connections in marmosets has not yet been examined. Our findings showing the significance of the Pf-Cd and CM-Pu systems in different learning processes of visual discrimination task suggest that these systems may interact with the corticostriatal inputs originating from distinct associative cortical areas, resulting in enhancement of the formation or switching of stimulus and response association. In this study, we did not test attentional set-shifting in the marmosets lacking Pf-Cd neurons, and it remains unclear whether the Pf-Cd system is also engaged in set-shifting of the visual discrimination task.

In rodents, the two major components of the ILN are the Pf and the central lateral nucleus (CL), which are localized in their caudal or rostral part, respectively. There are variations in synaptic structures and electrophysiological properties between these nuclei^49–52^. The thalamostriatal systems originating from the Pf and CL play essential roles in different learning processes of operant conditioning and sensory discrimination as well as in the flexible switching of learned behaviors^27,28,53–59^. Also homologically, the Pf of rodents projects mainly to the dorsolateral striatum, whereas the CL innervates the boundary between the dorsolateral and the dorsomedial striatum^60^. These lines of evidence, together with the present data obtained in common marmosets, suggest that the functional diversity between the two principal structures of the ILN may be closely related to their output projections toward distinct territories of the striatum in both rodents and nonhuman primates.

In addition, the thalamostriatal systems provide sparsely their axonal collaterals to some cortical areas in rodents, including the collaterals from the Pf to the frontal, motor, and insular cortical areas^61^ and those from the CL to the motor cortex^62^. Structural connectivity analysis of the human brain reports the distribution of axonal collaterals originating from the CM/Pf complex to the somatosensory cortical areas^63^, although the distribution of collaterals in the marmoset brain has not yet been fully investigated. These collaterals may affect some behavioral responses, such as the acquisition or switching of sensory cue-dependent decision making we tested herein. The physiological and behavioral significance of the collaterals from the ILN remains to be addressed in the future.

Human postmortem brain studies demonstrate 30–40% neuronal loss in the ILN complex in PD patients, in addition to substantia nigra dopamine neurons^22–24^. Monkeys chronically treated with MPTP also have a reduced number of synaptic terminals of the ILN neurons in the dorsal striatum^25,26^. The present results suggest that loss of the ILN neurons may be associated with some cognitive deficits in PD, such as the instrumental learning and flexible switching of learned behaviors, whereas motor skills and spontaneous locomotion appeared to be unaffected despite of the decreased neuronal number. Activation of thalamostriatal system has been shown to alleviate deficits in motor functions in PD model mice through striatal cholinergic interneurons^64,65^. In addition, blocking neuroinflammation in the Pf rescues cognitive impairment in PD model rats^66^. Thus, the CM/Pf complex may be a potential, effective target for therapeutic strategies for the symptoms of PD.

## Materials and Methods

### Experimental animals

All experiments were carried out in accordance with the guidelines established by Fukushima Medical University, Primate Research Institute of Kyoto University, Laboratory Animal Welfare established by CLEA Japan, Inc. (Tokyo, Japan), and the Central Research Center for Experimental Animals (Tokyo, Japan). All care and handling procedures were approved by their Institutional Animal Care and Use Committees. Sixteen common marmosets (*Callithrix jacchus*, 10 males and 6 females, aged 18–50 months, 285–384 g in body weight; CLEA Japan, Inc.) were named MAR0–MAR15 and used for the present study. The marmosets were housed individually in a home cage of 39 × 55 × 70 cm (W × D × H) in size, and maintained at 28 ± 1°C and 50 ± 10% humidity in a 12-h light/12-h dark cycle (7:00 to 19:00 for light). The experiments for two-choice visual discrimination and staircase foraging tasks were conducted between 9:00 and 17:00, and locomotion activity was measured for 12 h (7:00 to 19:00). They were fed 20–30 g New World Monkey Diet (CMS-1M; CLEA Japan, Inc.) once daily, and water was provided *ad libitum*. Four types of solid rewards (marshmallow, sponge cake, strawberry taste candy and orange taste candy) were provided during the behavioral test.

### Brain imaging

MR imaging and X-ray computed tomography (CT) were conducted as previously described^67^ with some modifications. Common marmosets were intramuscularly administered 0.1 mg/kg atropine sulfate and 12 mg/kg alfaxalone, and the anesthetized state was maintained under 1–2% isoflurane inhalation. The marmosets were placed in an acrylic imaging cradle (Takashima Seisakusho Ltd., Tokyo, Japan), and an acrylic head holder was attached to their heads, inserting ear bars into the external auditory canals. Pulse oxygen and skin and rectal temperature were monitored regularly during the imaging. MR image data were obtained using the 7.0 T Biospec 70/16 MR Image Scanner System (Bruker BioSpin GmbH; Ettlingen, Germany) equipped with actively shielded gradients at a maximum strength of 700 mT/m and an imaging coil (inner diameter 60 mm; Bruker Biospin GmbH). T2-weighted images were acquired using a rapid acquisition with relaxation enhancement sequence (RARE) with the following parameters: repetition time 4,500 ms, echo time 20 ms, field of view (FOV) 48 × 48 mm, matrix 240 × 240, slice thickness 0.35 mm, RARE factor 4, number of average 11, and scan time 42 min. Then CT image data were obtained at the same scanning position as the MR imaging by using a cone-beam CT system (Cosmo Scan Fx; Rigaku Corp., Tokyo, Japan), which was operated under the following conditions: tube voltage 90 kV, tube current 88 μA, exposure time 2 min, FOV 61.44 × 61.44 × 61.44 mm, and voxel size 120 μm^3^. The skull was semi-automatically extracted from CT data using the “Segmentation Editor” in Amira software version 7.0 (Visage Imaging, Inc., San Diego, CA, USA). The MR and CT images were manually overlaid based on the position of ear bars as the stereotaxic landmark. CT images were resliced to resample isotropic MR images (0.2 mm^3^). MR and CT images used for injection coordinates were viewed with PMOD image analysis software version 3.7 (PMOD Technology Ltd., Fällanden, Switzerland). Injection coordinates were calculated using Adobe Photoshop.

### Viral vector preparation

NeuRet vectors were prepared and purified as previously described^29^. The transfer plasmids (pCL20c-MSCV-GFP, pCL20c-MSCV-cgfTagRFP and pCL20c-MSCV-IL2Rα/GFP) contained the cDNAs encoding GFP, a mutant form of RFP (cgfTagRFP)^68^, and human IL2Rα fused to GFP^69^, respectively, downstream of the murine stem cell virus promoter. HEK293T cells were transfected with the transfer, envelope containing fusion glycoprotein type E (FuG-E)^30^ cDNA and packaging plasmids using the calcium phosphate precipitation method. After viral collection and purification, quantitative PCR was carried using the StepOne Real-Time PCR System (Applied Biosystems, Tokyo, Japan) with the Lenti-X qRT-PCR Titration Kit (Takara Bio Inc., Kusatsu, Japan).

### Intracranial surgery

Marmosets were placed in a stereotaxic instrument (SR-6C-HT; Narishige, Tokyo, Japan) under anesthesia as described above. The marmosets were administered viccillin (2.5–10 mg/kg, intramuscular injection) as an antibiotic drug and meloxicam (0.5 mg/kg, subcutaneously) as an anti-inflammatory drug, together with Ringer’s solution (5 mL, subcutaneously) containing DL-methionine (4.5 mg), thiamine chloride hydrochloride (300 μg), sodium riboflavin phosphate (60 μg), pyridoxine hydrochloride (150 μg), nicotinic acid amide (750 μg) and sodium L-ascorbate (3.0 mg). After hair removal and sterilization with povidone iodine, burr holes were opened on the skull for injections of the viral vectors and ITX/PBS solution. The vectors were introduced into the head part of the Cd (1.0 μL/site, 6 sites/hemisphere) or the anteroposterior middle part of the Pu (1.0 μL/site, 4 sites/hemisphere), and ITX solution (25 ng/µL in PBS containing 0.1 mg/mL monkey serum albumin) or PBS was injected into the Pf (0.5 μL/site, 1 site/hemisphere) or CM (0.5 μL/site, 2 sites/hemisphere) through a glass microinjection capillary connected to a microinfusion pump (ESP-32; Eicom, Kyoto, Japan). The glass microinjection capillary was inserted into the brain, moved slowly to 0.2 mm beyond the target, and then pulled up to the target position. For the retrograde labeling of neural pathways, the anteroposterior, mediolateral, and dorsoventral coordinates (mm) from the interaural line and brain surface were 12.5/2.8/5.8 (site 1), 12.5/2.8/5.3 (site 2), 10.5/3.3/5.7 (site 3), 10.5/3.3/5.2 (site 4), 8.5/3.2/5.9 (site 5), and 8.5/3.2/5.2 (site 6) for Cd injection; and 10.0/6.2/6.7 (site 1), 10.0/6.2/5.7 (site 2), 8.5/6.5/6.9 (site 3), and 8.5/6.5/5.9 (site 4) for Pu injection. For the selective targeting of neural pathways, the coordinates in each animal are summarized in Supplementary Table 1 (for Cd/Pf injections) and 2 (for Pu/CM injections). Injection was performed at a constant flow rate of 0.2 μL/min for vectors and 0.1 μL/min for ITX/PBS solution.

### Histological analysis

The perfusion fixation was conducted under anesthesia with a mixture of medetomidine (0.04 mg/kg), midazolam (0.4 mg/kg), and butorphanol (0.4 mg/kg) for induction and isoflurane (4%) for terminal care. Marmosets were perfused transcardially with PBS, followed by fixation in 4% paraformaldehyde in 0.1 M PBS (pH 7.4). The brains were removed from the skull, postfixed for 1–2 days, and saturated with 15% and 30% sucrose in PBS at 4°C. Fixed brains were cut into sections (30 μm thick) through a coronal plane with a cryostat. For immunohistochemistry, sections were incubated overnight at 4°C with anti-GFP antibody (rabbit, 1:2000, #GFP-Rb-Af2020; Nittobo Medical Co., Ltd., Tokyo, Japan), anti-RFP (guinea pig, 1:1000, Nittobo Medical), anti-NeuN antibody (mouse, 1:400, #MAB377; Merck Millipore, Burlington, MA, USA), or anti-IL2Rα antibody (goat, 1:400, #I6152; Merck Millipore). Then the sections were incubated with species-specific IgG antibodies conjugated to biotin (for rabbit, 1:500, #711-065-152; Jackson ImmunoResearch Labs, West Grove, PA, USA or for goat, 1:500, #711-065-147; Jackson ImmunoResearch Labs), Cy3 (for guinea pig, 1:500, #706-165-148; Jackson ImmunoResearch Labs), Alexa 488 (for rabbit, 1:500, #A-21206; Thermo Fisher Scientific, Waltham, MA, USA) or Alexa 647 (for mouse, 1:500, #A-31571; Thermo Fisher Scientific) for 2 h at room temperature. The sections were treated with an avidin-biotin-peroxidase complex kit (VECTASTAIN Elite ABC kit; Vector Laboratories Inc., Newark, CA, USA). To visualize gene expression, sections were reacted with 0.05 M Tris-HCl buffer (pH 7.6) containing 0.05% 3,3’-diaminobenzidine tetrahydrochloride (DAB; DOJINDO, Kumamoto, Japan) and 0.3% H_2_O_2_ (FUJIFILM WAKO Pure Chemical, Osaka, Japan) for 5–10 min. These sections were mounted on gelatin-coated glass slides and coverslipped. The images were acquired using a fluorescent microscope (BZ-X810; Keyence, Osaka, Japan).

### Cell counts

The number of immunopositive cells in each of 10 sections stained at 100-µm intervals for the Pf or at 150-µm intervals for the CM were counted, and the values for individual animals were averaged.

### Behavioral analysis

Two-choice visual discrimination and staircase foraging tasks were executed in an experimental cage of 39 × 55 × 70 cm in size, which was modified from the home cage to equip the apparatus used for the respective tasks. Each animal was transferred individually from the home cage to the experimental cage by using a small transport chamber of 135 × 235 × 150 mm in size (JIC Co., Ltd., Kawasaki, Japan).

### Two-choice visual discrimination task

The visual discrimination task was conducted according to previous studies^34,35^ with some modifications. The apparatus consisted of a 15-inch screen with a black background installed to a computer, a USB-driven feeder connected to a food tray below the screen, and a tone generator, attaching to the front panel of the experimental cage. During the habituation phase, each marmoset was trained to touch a single visual image on the screen. Initially, a colored square image (red, blue, or yellow) of a large size (75 × 75 mm) was presented randomly at the center of the screen. When the marmoset touched the image, it disappeared with presentation of a tone (4 kHz, 100 ms), and a solid reward was delivered into the food tray through the feeder. Each daily session ended when 60 trials were completed or when 60 min had elapsed with the intertrial interval (ITI) of 3 s. The sessions continued for 2 days. Next, the size of the colored square images was reduced to 30 × 30 mm and presented at the center of the screen, the sessions were carried out as described above. The image was then presented randomly at different positions on the screen, and the sessions continued for 2 days.

For the acquisition phase, the initiation of each trial was signaled by presentation of a red square image (30 × 30 mm) at the center of the screen. After a 1-s interval, a pair of square graphic images (see Supplementary Fig. 1) was presented simultaneously on the left and right sides of the screen. The left-right positions were randomly changed and counter-balanced in a session. One of the images (correct image) was associated with a reward and the other (error image) was not. When the marmoset touched either of the images, both disappeared. When the marmoset touched the correct image, a solid reward was delivered with the tone presentation. Each session ended when 50 trials were completed or when 60 min had elapsed. The correct and error responses were followed by 3- and 5-s ITIs, respectively. The sessions continued for 10 days. When a session ended less than 50 trials in a 60-min period, the session was aborted. The numbers of trials, which reached a criterion of 75% correction, were calculated to evaluate the acquisition of visual discrimination.

For the reversal phase, the behavioral task was performed as described above except for changing the correct image to another one in the pair of images used for the acquisition phase. Error responses in the reversal phase were classified into perseverative and regressive errors, which were defined as the errors that occur until an animal first selects the new correct image and that occur afterwards, respectively^36,37^.

### Staircase foraging task

The foraging task was conducted according to previous reports^38,39^ with some modifications. The apparatus consisted of two 5-step staircases on the left and right side with an acrylic plate slit at the center, attaching to the front panel of the experimental cage. The marmosets had to reach and retrieve one of the solid rewards through the plate slit by using either right or left arm opposite to the side of the stairs. The task was performed 4 trials in a daily session on each side (total of 8 trials/session) and each trial was for 5 min. The session continued for consecutive 10 days. The correct ratio was calculated as the percentage of the number of successful trials divided by the total number of trials. All sessions were video-recorded, and the correct and error responses were determined by video judgment.

### Locomotion activity measurement

Spontaneous locomotion activity was counted in the home cage by using a miniatured accelerometer with 128 kilobytes of memory in a cylindrical housing (diameter 21 mm, height 8 mm, weight 4.4 g; Actiwatch-Mini, CamNtech Ltd., Cambridge, UK)^40,41^. Spontaneous activity was binned and recorded from all animals per day. The accelerometer was programmed to record the data for every 60 s, which were calculated as the total amount of activity for 12 h.

### Statistical analyses

All data are presented as mean ± standard error of the mean (SEM). For multiple comparisons, two-way ANOVA was applied followed by post hoc Bonferroni test. The unpaired Student’s *t*-test was also used. All statistical details of the experiments can be found in the Results section. Differences were considered statistically significant at *p* < 0.05.

### Reporting summary

Further information on the research design is available in the Nature Portfolio Reporting Summary linked to this article.

### Data availability

The authors declare that all data supporting the findings of this study are available within the article and its supplementary information file, and are available from the corresponding author upon request without restrictions. Source data are deposited on Mendeley data (https://data.mendeley.com/datasets/c5b9tbs3hr/1).

## Acknowledgments

This work was supported by a grant-in-aid from the Japan Agency for Medical Research and Development (No. 22dm0207113h0002 to K.K. and 22dm0207077 to M.T.); a grant-in-aid for Scientific Research on Transformative Research Areas (A) Adaptive Circuit Census (No. 21H05244 to K.K.) and Biology of Behavior Change (No. 22H05157 to K.I.); and a grant-in-aid for Scientific Research (C) (No. MO20K06912 to S.K.) from the Japan Society for the Promotion of Science from the Ministry of Education, Science, Sports, and Culture of Japan. We are grateful to K. Fukasawa, K. Kagiyama, H. Hibino, and S. Maeda for their invaluable help with the animal treatment and care; to H. Hashimoto, M. Kikuchi, and Y. Nakazato for their technical support for vector preparation; and to T. Kobayashi for her helpful illustrations.

## Author contributions

S.K., M.S., K.N., M.T., and K.K. conceived the study, designed the experiments, and directed the project. S.K. designed the vector expression strategy and generated the vectors. M.S., M.Y., K.I., and M.W. performed intracranial injections and histological examinations. S.K., and K.K. wrote the paper. All authors discussed the results and implications, and commented on the manuscript at all stages.

