## Supplemental information for "Distinct roles of two thalamostriatal systems in learning processes of visual discrimination in common marmosets"

**Supplementary Table 1 | Coordinates of injection of vector (Cd), ITX and PBS (Pf) in each marmoset with functional manipulation of the Pf-Cd pathway.**

| Injection | PBS | | | | | | | | | ITX | | | | | | | | | | | |
|---|---|---|---|---|---|---|---|---|---|---|---|---|---|---|---|---|---|---|---|---|---|
| Animal | MAR-1 | | | MAR-2 | | | MAR-3 | | | MAR-4 | | | MAR-5 | | | MAR-6 | | | MAR-7 | | |
| Axis | AP | ML | DV | AP | ML | DV | AP | ML | DV | AP | ML | DV | AP | ML | DV | AP | ML | DV | AP | ML | DV |
| Cd1 | 12.6 | -2.8 | 5.8 | 12.4 | -2.7 | 5.8 | 13.2 | -2.7 | 6.4 | 12.6 | -3.1 | 6.3 | 12.8 | -3.0 | 6.4 | 13.6 | -2.8 | 5.8 | 10.3 | -2.5 | 5.8 |
| Cd2 | 12.6 | -2.8 | 5.3 | 12.4 | -2.7 | 5.3 | 13.2 | -2.7 | 5.9 | 12.6 | -3.1 | 5.8 | 12.8 | -3.0 | 5.9 | 13.6 | -2.8 | 5.3 | 10.3 | -2.5 | 5.3 |
| Cd3 | 12.6 | 2.8 | 5.8 | 12.4 | 2.9 | 6.0 | 13.2 | 2.8 | 6.4 | 12.6 | 3.0 | 6.3 | 12.8 | 2.9 | 6.5 | 13.6 | 3.3 | 5.7 | 10.3 | 2.5 | 5.8 |
| Cd4 | 12.6 | 2.8 | 5.3 | 12.4 | 2.9 | 5.5 | 13.2 | 2.8 | 5.9 | 12.6 | 3.0 | 5.8 | 12.8 | 2.9 | 6.0 | 13.6 | 3.3 | 5.2 | 10.3 | 2.5 | 5.3 |
| Cd5 | 11.0 | -3.0 | 5.6 | 10.2 | -3.0 | 5.8 | 11.0 | -3.1 | 6.4 | 10.4 | -3.0 | 6.7 | 10.6 | -2.9 | 6.5 | 11.6 | -3.2 | 5.9 | 8.8 | -2.6 | 5.7 |
| Cd6 | 11.0 | -3.0 | 5.1 | 10.2 | -3.0 | 5.3 | 11.0 | -3.1 | 5.9 | 10.4 | -3.0 | 6.2 | 10.6 | -2.9 | 6.0 | 11.6 | -3.2 | 5.2 | 8.8 | -2.6 | 5.2 |
| Cd7 | 11.0 | 3.0 | 5.6 | 10.2 | 3.1 | 6.0 | 11.0 | 2.8 | 6.7 | 10.4 | 2.9 | 6.8 | 10.6 | 3.0 | 6.5 | 11.6 | 2.8 | 5.8 | 8.8 | 2.6 | 5.7 |
| Cd8 | 11.0 | 3.0 | 5.1 | 10.2 | 3.1 | 5.5 | 11.0 | 2.8 | 6.2 | 10.4 | 2.9 | 6.3 | 10.6 | 3.0 | 6.0 | 11.6 | 2.8 | 5.3 | 8.8 | 2.6 | 5.2 |
| Cd9 | 8.6 | -3.1 | 5.6 | 8.0 | -3.3 | 5.6 | 8.8 | -3.2 | 6.4 | 8.2 | -3.3 | 6.3 | 8.4 | -3.0 | 6.4 | 9.6 | -3.3 | 5.7 | 6.6 | -3.0 | 5.3 |
| Cd10 | 8.6 | -3.1 | 5.1 | 8.0 | -3.3 | 5.1 | 8.8 | -3.2 | 5.9 | 8.2 | -3.3 | 5.8 | 8.4 | -3.0 | 5.9 | 9.6 | -3.3 | 5.2 | 6.6 | -3.0 | 4.8 |
| Cd11 | 8.6 | 3.1 | 5.6 | 8.0 | 3.2 | 5.8 | 8.8 | 2.9 | 6.3 | 8.2 | 3.1 | 6.3 | 8.4 | 3.1 | 6.4 | 9.6 | 3.2 | 5.9 | 6.6 | 3.0 | 5.3 |
| Cd12 | 8.6 | 3.1 | 5.1 | 8.0 | 3.2 | 5.3 | 8.8 | 2.9 | 5.8 | 8.2 | 3.1 | 5.8 | 8.4 | 3.1 | 5.9 | 9.6 | 3.2 | 5.2 | 6.6 | 3.0 | 4.8 |
| Pf1 | 4.4 | -0.9 | 10.0 | 3.6 | -1.0 | 10.3 | 4.4 | -0.8 | 9.8 | 3.6 | -0.9 | 10.5 | 3.8 | -0.9 | 10.0 | 5.4 | -1.0 | 10.1 | 2.8 | -0.8 | 9.5 |
| Pf2 | 4.4 | 0.9 | 10.0 | 3.6 | 0.8 | 10.3 | 4.4 | 0.7 | 10.0 | 3.6 | 0.8 | 10.5 | 3.8 | 0.9 | 10.2 | 5.4 | 0.7 | 10.2 | 2.8 | 0.8 | 9.5 |

**Supplementary Table 2 | Coordinates of injection of vector (Pu), ITX and PBS (CM) in each marmoset with functional manipulation of the CM-Pu pathway.**

| Injection | PBS | | | | | | | | | | | | ITX | | | | | | | | | | | |
|---|---|---|---|---|---|---|---|---|---|---|---|---|---|---|---|---|---|---|---|---|---|---|---|---|
| Animal | MAR-8 | | | MAR-9 | | | MAR-10 | | | MAR-11 | | | MAR-12 | | | MAR-13 | | | MAR-14 | | | MAR-15 | | |
| Axis | AP | ML | DV | AP | ML | DV | AP | ML | DV | AP | ML | DV | AP | ML | DV | AP | ML | DV | AP | ML | DV | AP | ML | DV |
| Pu1 | 8.6 | -5.4 | 7.0 | 11.0 | -5.4 | 7.2 | 10.8 | -5.9 | 6.8 | 10.4 | -5.6 | 7.5 | 11.2 | -5.4 | 7.8 | 9.8 | -5.3 | 7.4 | 10.6 | -5.9 | 6.9 | 9.8 | -5.5 | 7.8 |
| Pu2 | 8.6 | -5.4 | 6.2 | 11.0 | -5.4 | 6.4 | 10.8 | -5.9 | 6.1 | 10.4 | -5.6 | 6.7 | 11.2 | -5.4 | 7.3 | 9.8 | -5.3 | 6.6 | 10.6 | -5.9 | 6.2 | 9.8 | -5.5 | 7.0 |
| Pu3 | 8.6 | 5.1 | 7.0 | 11.0 | 5.4 | 7.6 | 10.8 | 5.9 | 7.0 | 10.4 | 5.6 | 7.4 | 11.2 | 5.3 | 7.7 | 9.8 | 5.2 | 7.4 | 10.6 | 5.6 | 6.9 | 9.8 | 5.6 | 7.8 |
| Pu4 | 8.6 | 5.1 | 6.2 | 11.0 | 5.4 | 6.8 | 10.8 | 5.9 | 6.3 | 10.4 | 5.6 | 6.6 | 11.2 | 5.3 | 7.2 | 9.8 | 5.2 | 6.6 | 10.6 | 5.6 | 6.2 | 9.8 | 5.6 | 7.0 |
| Pu5 | 7.2 | -6.0 | 6.7 | 9.6 | -6.4 | 6.9 | 9.4 | -6.3 | 6.5 | 9.0 | -6.2 | 7.0 | 9.6 | -6.2 | 7.2 | 8.4 | -6.1 | 6.6 | 9.2 | -6.5 | 6.4 | 8.4 | -6.1 | 7.3 |
| Pu6 | 7.2 | -6.0 | 5.9 | 9.6 | -6.4 | 6.1 | 9.4 | -6.3 | 5.8 | 9.0 | -6.2 | 6.2 | 9.6 | -6.2 | 6.7 | 8.4 | -6.1 | 5.8 | 9.2 | -6.5 | 5.7 | 8.4 | -6.1 | 6.5 |
| Pu7 | 7.2 | 5.5 | 6.8 | 9.6 | 6.0 | 7.3 | 9.4 | 6.2 | 6.7 | 9.0 | 6.2 | 7.0 | 9.6 | 6.0 | 7.2 | 8.4 | 6.0 | 6.8 | 9.2 | 6.4 | 6.7 | 8.4 | 6.2 | 7.4 |
| Pu8 | 7.2 | 5.5 | 6.0 | 9.6 | 6.0 | 6.5 | 9.4 | 6.2 | 6.0 | 9.0 | 6.2 | 6.2 | 9.6 | 6.0 | 6.7 | 8.4 | 6.0 | 6.0 | 9.2 | 6.4 | 6.0 | 8.4 | 6.2 | 6.6 |
| CM1 | 3.6 | -1.6 | 9.2 | 5.6 | -1.7 | 9.5 | 5.4 | -1.5 | 9.5 | 4.9 | -1.7 | 10.3 | 5.8 | -1.2 | 9.9 | 5.0 | -1.6 | 9.8 | 5.2 | -1.8 | 9.5 | 4.4 | -1.7 | 10.5 |
| CM2 | 3.6 | 1.1 | 9.1 | 5.6 | 1.5 | 9.5 | 5.4 | 1.3 | 9.5 | 4.9 | 1.7 | 10.4 | 5.8 | 1.4 | 10.2 | 5.0 | 1.4 | 10.0 | 5.2 | 1.6 | 9.5 | 4.4 | 1.7 | 10.5 |
| CM3 | 3.2 | -1.9 | 9.3 | 5.0 | -1.7 | 10.1 | 4.9 | -1.6 | 9.5 | 4.4 | -1.7 | 10.2 | 5.2 | -1.4 | 10.5 | 4.4 | -1.6 | 10.2 | 4.7 | -1.8 | 9.6 | 3.9 | -1.7 | 10.5 |
| CM4 | 3.2 | 1.3 | 9.0 | 5.0 | 1.6 | 10.1 | 4.9 | 1.4 | 9.5 | 4.4 | 1.8 | 10.3 | 5.2 | 1.4 | 10.8 | 4.4 | 1.4 | 10.4 | 4.7 | 1.6 | 9.6 | 3.9 | 1.7 | 10.5 |

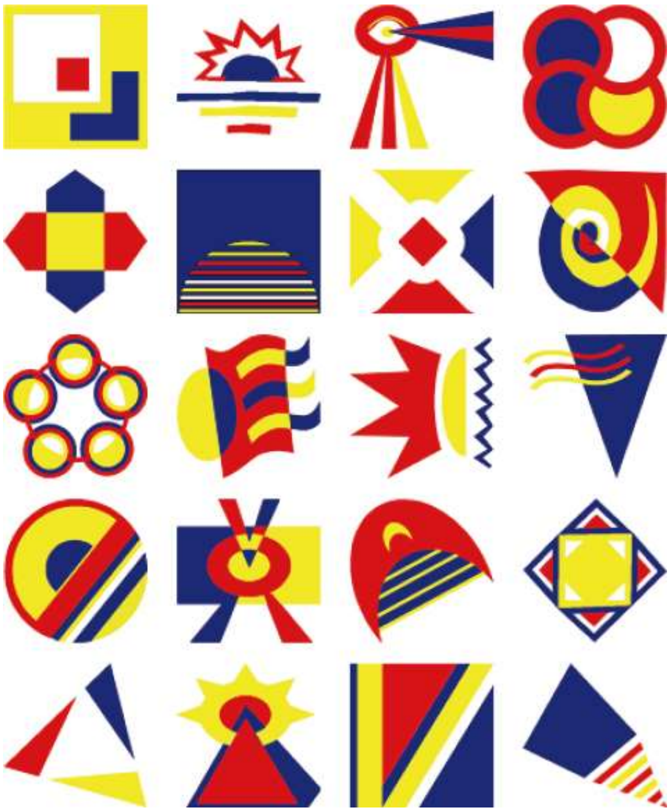

**Supplementary Fig. 1 | Visual images used for a two-choice visual discrimination task.** A pair of images was randomly selected from the above 20 different images for each animal.

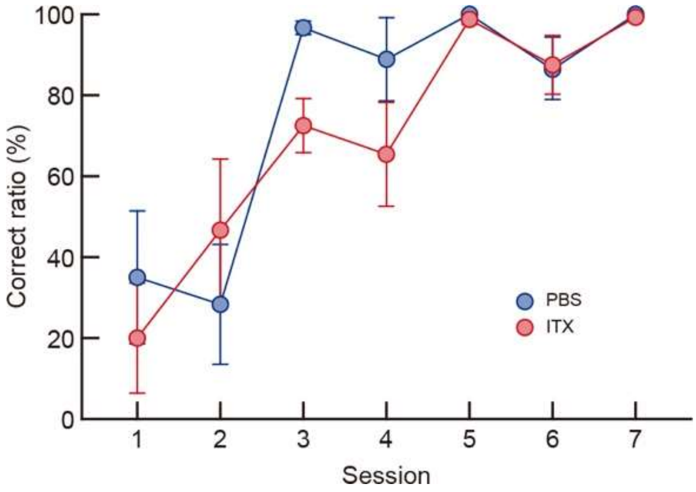


**Supplementary Fig. 2 | Learning curve of a simple touch during the habituation phase of common marmosets lacking Pf-Cd neurons.** The correct ratio in each session was plotted in accordance with the progress of training (two-way ANOVA: group, $F_{(1, 5)} = 0.991$, $p = 0.365$; session, $F_{(6, 30)} = 16.623$, $p < 0.001$; interaction, $F_{(6, 30)} = 1.147$, $p = 0.360$). $n = 4$ for the ITX-injected group, and $n = 3$ for the PBS-injected group. Data are presented as the mean ± SEM.

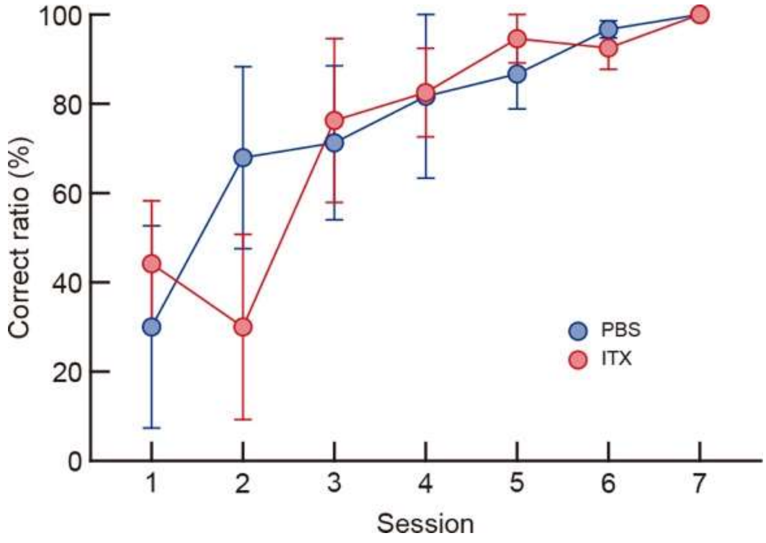


**Supplementary Fig. 3 | Learning curve of a simple touch during the habituation phase of common marmosets lacking CM-Pu neurons.** The correct ratio in each session was plotted in accordance with the progress of training (two-way ANOVA: group, $F_{(1, 6)} = 0.024$, $p = 0.883$; session, $F_{(6, 36)} = 8.936$, $p < 0.001$; interaction, $F_{(6, 36)} = 1.121$, $p = 0.370$). $n = 4$ for each group. Data are presented as the mean ± SEM.
